# Targeting Diacylglycerol Lipase to Reduce Alcohol Consumption

**DOI:** 10.1101/2021.02.16.431429

**Authors:** Gaurav Bedse, Nathan D. Winters, Anastasia Astafyev, Toni A. Patrick, Vikrant R. Mahajan, Md. Jashim Uddin, Samuel W. Centanni, David C. Samuels, Lawrence J. Marnett, Danny G. Winder, Sachin Patel

## Abstract

Alcohol use disorder (AUD) is associated with substantial morbidity, mortality, and societal cost, and pharmacological treatment options for AUD are limited. The endogenous cannabinoid (eCB) signaling system is critically involved in reward processing and alcohol intake is positively correlated with release of the eCB ligand 2-Arachidonoylglycerol (2-AG) within reward neurocircuitry. Here we show that genetic and pharmacological inhibition of diacylglycerol lipase (DAGL), the rate limiting enzyme in the synthesis of 2-AG, reduces alcohol consumption in a variety of preclinical models ranging from a voluntary free-access model to aversion resistant-drinking, and dependence-like drinking induced via chronic intermittent ethanol vapor exposure in mice. DAGL inhibition also prevented ethanol-induced suppression of GABAergic transmission onto midbrain dopamine neurons, providing mechanistic insight into how DAGL inhibition could affect alcohol reward. Lastly, DAGL inhibition during either chronic alcohol consumption or protracted withdrawal was devoid of anxiogenic and depressive-like behavioral effects. These data suggest reducing 2-AG signaling via inhibition of DAGL could represent a novel approach to reduce alcohol consumption across the spectrum of AUD severity.

## INTRODUCTION

Alcohol use disorder (AUD) is a chronic relapsing substance abuse disorder that exhibits a lifetime prevalence of approximately 30% in the United States (1). Like disordered use of other drugs, AUD can be conceptualized as a chronic disease quintessentially characterized by lack of control over alcohol drinking behaviors, despite adverse consequences. Patients with AUD exhibit cravings and a compulsive drive to seek and use alcohol during abstinence, and may exhibit tolerance and withdrawal symptoms often leading to relapse (2). Societal costs related to problematic drinking led to an estimated economic burden of $249 billion in the United States in 2010 (3), emphasizing AUD as a critical public health concern. Current treatment approaches for AUD consist of limited pharmacological treatments combined with various forms of group and individual psychotherapy, often and most effectively in conjunction (2). Despite this, efficacy is limited and relapse rates in AUD are high (2, 4), highlighting the need for the elucidation of novel and effective pharmacological targets for the advancement of AUD therapeutics development.

A growing body of work has implicated the endocannabinoid (eCB) system as a critical modulator of the effects of ethanol (see (5) and (6) for review), and the eCB system is heavily implicated in a variety of processes relevant to distinct stages of AUD including reward processing, stress-reactivity, and affect modulation (7–9), suggesting this system may serve as a promising target for novel AUD therapeutics. The eCB system is a retrograde neuromodulator system wherein the lipid-derived eCB ligands, 2-arachidonoylglycerol (2-AG) and anandamide (AEA), are synthesized and released from the postsynaptic compartments of neurons and activate presynaptic cannabinoid-1 (CB_1_) receptors to decrease neurotransmitter release probability. EtOH mobilizes the major brain eCB 2-AG in the nucleus accumbens (10, 11) and in some neuronal culture models (12, 13), and inhibition of CB_1_ receptors attenuates EtOH self-administration in rodents (14, 15) and diminishes ethanol-stimulated enhancements of dopamine neuron activity in midbrain reward circuits (16), collectively suggesting a necessity for intact eCB signaling for driving alcohol-seeking behaviors.

Inhibition of eCB signaling is posited as a potentially effective strategy to treat AUD, and a CB_1_ receptor inverse agonist (Rimonabant) has been explored in clinical trials for AUD (17) as well as other addiction-related disorders (18, 19), but this compound was removed from the European Union market due to severe neuropsychiatric side effects (20). Given that EtOH mobilizes 2-AG in the reward circuitry in a manner correlated with EtOH intake (10, 11), we hypothesized targeted inhibition of 2-AG signaling (thereby leaving AEA-CB_1_ signaling intact) could be an effective approach to reducing EtOH drinking with a lower adverse effect liability. Here we implemented genetic and pharmacological strategies to test the hypothesis that inhibiting diacylglycerol lipase (DAGL), the rate-limiting enzyme in 2-AG synthesis, reduces EtOH drinking behaviors without facilitating the development of anxiety- and depression-like behaviors after chronic EtOH drinking. We found that genetic deletion of DAGLα, the primary DAGL in the adult brain (21), or the pharmacological DAGL inhibitor DO34 (22) decreased EtOH consumption across a range of clinically-relevant models and does not exacerbate negative affective behaviors in chronically drinking mice or during protracted abstinence. Collectively, our data suggest that DAGL inhibition could be a promising and broadly applicable therapeutic strategy to reduce alcohol drinking across the spectrum of AUD.

## RESULTS

### Genetic and pharmacological inhibition of DAGL decreases voluntary EtOH drinking

Having previously established that DAGLα genetic deletion results in reduced brain 2-AG levels (9), we evaluated the effects of DAGLα deletion on EtOH drinking behavior. Separate cohorts of male and female DAGLα^-/-^ mice and WT littermates were exposed to two-bottle choice (2BC) EtOH drinking paradigm (**Fig. 1a**). Both male and female DAGLα^-/-^ mice exhibited lower EtOH preference (**Fig. 1b, d**) and consumption (**Fig. 1c, e**) relative to their WT littermates. DAGLα^-/-^ mice exhibited no difference in total fluid consumption (**Fig. S2a-b**). We next evaluated the effects of pharmacological inhibition of DAGL on alcohol drinking by using the DAGL inhibitor DO34 (22). Previously, we have demonstrated that DO34 (50 mg/kg) significantly decreases brain 2-AG levels after 2 hours of administration (9). Separate cohorts of male and female C57BL/6J mice were exposed to the 2BC EtOH drinking paradigm. After 6 weeks of stable drinking, DO34 (50 mg/kg) was administered for three consecutive days. DO34 significantly reduced EtOH preference (**Fig. 1g, i**) and consumption (**Fig. 1h, j**). DO34 did not significantly alter total fluid consumption in either sex (**Fig. S2c-d**). Vehicle control injections had no effect on EtOH preference (**Fig. S3a, d**) or consumption (**Fig. S3b, e**). DO34 had no effect on body weight in female mice (**Fig. S3c**) but caused a small but significant decrease in body weight in males that recovered upon cessation of drug treatment (**Fig. S3f**). Due to their higher relative EtOH preference and consumption levels, female mice were used for all drinking and behavioral experiments hereafter.

**Figure 1:**
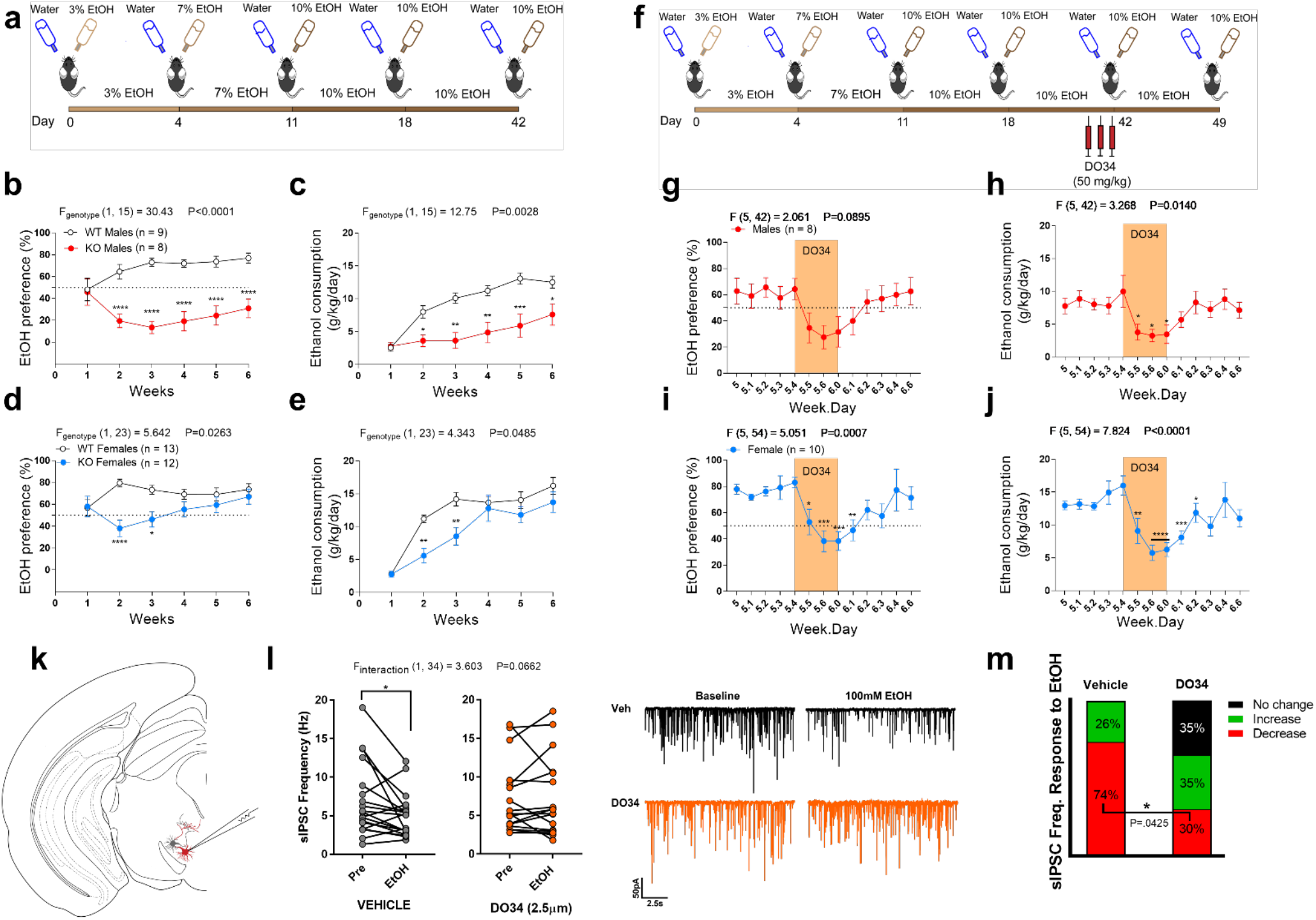
Inhibition of DAGL signaling reduces 2BC alcohol drinking and prevents alcohol suppression of VTA GABA transmission. **(a)** Schematic of 2BC EtOH drinking paradigm for DAGLα^-/-^mice. **(b)** Male DAGLα^-/-^ mice exhibited lower EtOH preference and **(c)** consumption relative to WT males. **(d)** Female DAGLα^-/-^ mice initially exhibited lower preference and **(e)** consumption that reached WT levels at week 4. **(f)** Schematic of 2BC EtOH drinking for WT mice receiving DO34 treatment. **(g)** Male WT mice exhibited a non-significant reduction in preference (Week.Day 5.6, P=.052) and a **(h)** significant reduction in EtOH consumption in response to DO34 treatment. **(i)** Female WT mice exhibited a significant reduction in EtOH preference and **(j)** consumption in response to DO34 treatment. **(k)** Schematic depicting recording strategy used to identify putative dopamine neurons exhibiting red fluorescence in the posterior VTA. **(l, left)** Bath application of 100mM EtOH reduced sIPSC frequency in vehicle-treated but not DO34-treated slices. **(l, right)** Example traces of sIPSC response to 100mM bath application in vehicle-(black) or DO34-treated (orange) slices. **(m)** Relative proportions of sIPSC frequency response to 100mM EtOH bath application, defined as ≥10% deviation from baseline. DO34 treatment significantly reduced the proportion of cells that exhibited a reduction in sIPSC frequency. Vehicle, n = 19 cells; DO34, n = 17 cells; from 13 mice in **l-m. (b-e, l)** Data analyzed by repeated measures two-way ANOVA followed by a Holm-Sidak test for multiple comparisons between genotypes **(b-e)** or baseline and EtOH treatment **(l)** or **(g-j)** one-way ANOVA on time points 5.4 – 6.2 (to include baseline, drug treatment, and two recovery points) followed by a Holm-Sidak test for multiple comparisons to baseline control. **(m)** Data analyzed by Fisher’s exact test for proportion of cells exhibiting reduced sIPSC frequency after EtOH application. Sample size *n, P* and *F* values for main effects of genotype or drug treatment reported in **b-e** and **g-j**, *P* and *F* value for EtOH x drug interaction reported in **l**. Significance for post-hoc multiple comparisons reported on graphs (* P<.05, * * P<0.01, * * * P<0.001, * * * * P<0.0001). Data in drinking experiments are mean ± SEM.

Although our convergent genetic and pharmacological studies indicate a role for DAGL in alcohol consumption, DO34 does exhibits some activity at other targets, such as alpha/beta hydrolase domain-containing 2 and 6 (ABHD2/6), platelet-activating factor acetylhydrolase 2 (PAFAH2), carboxylesterase 1C (CES1C), and phospholipase A2 group 7 (PLA2G7) (22). To confirm that the DO34 effects on EtOH consumption and preference are specific to DAGL inhibition, we utilized the structural analog DO53, which retains the off-target activity profile of DO34 but does not inhibit DAGL (22). A cohort of female mice stably drinking 10% EtOH on our 2BC paradigm were administered DO53 (50 mg/kg) for 5 consecutive days. EtOH consumption was transiently reduced by DO53 treatment (**Fig. S4a**), however, this was paralleled by a non-specific decrease in total fluid consumption (**Fig. S4c**). Accordingly, there was no effect on EtOH preference (**Fig. S4b**) and no change in body weight (**Fig. S4d**) during the 5-day DO53 treatment. These effects are in contrast with the persistent decrease in EtOH preference and consumption (**Fig. 1i-j**) and unchanged fluid consumption (**Fig. S2c-d**) observed after DO34 treatment. Combined with our genetic studies using DAGLα^-/-^ mice, these control data suggest the effects of DO34 were mediated through DAGL inhibition, rather than off-target mechanisms.

### DO34 Blocks EtOH-Driven Disinhibition of Posterior VTA Dopamine Neurons

The negative effect of DO34 on alcohol consumption suggests a role for 2-AG mobilization in driving alcohol-seeking. EtOH stimulates the activity of VTA dopamine neurons in a manner that requires eCB signaling (16). Accordingly, it has been previously reported that EtOH reduces inhibitory GABA transmission onto putative dopamine neurons in the posterior VTA (23), and VTA GABA transmission is regulated by eCB signaling (24–27). Furthermore, it has been reported that DAGL lipase signaling disinhibits dopamine neurons by suppressing GABA release following chronic nicotine exposure (24), suggesting EtOH may stimulate a similar eCB-mediated disinhibitory mechanism on these cells. To test whether DAGL regulates alcohol-induced inhibition of VTA GABAergic transmission, we utilized tyrosine hydroxylase (TH)::Ai14 mice that express Td-tomato in TH+ cells to conduct fluorescence-assisted electrophysiological recordings of spontaneous inhibitory postsynaptic currents (sIPSCs) from putative dopamine (TH+) neurons in posterior midbrain slices (**Fig. 1k**). Acute bath application of 100mM EtOH decreased the frequency of sIPSCs onto the majority of cells recorded, as previously reported (23) (**Fig. 1l**). Incubation of slices in 2.5µM DO34 abolished this effect (**Fig. 1l**) and decreased the proportion of cells that exhibited a reduction in sIPSC frequency in response to EtOH application (**Fig. 1m**). Additionally, EtOH induced a small but significant reduction in sIPSC amplitude in DO34-treated slices (**Fig. S5**). These data suggest DO34 could decrease positive-reinforcement driven voluntary EtOH consumption via preventing EtOH-induced reductions in VTA GABA transmission and subsequent increases in VTA dopamine neuron activity.

### Effects of 2-AG modulation are specific to EtOH and not bidirectional

To confirm that DO34 effects are specific to EtOH and not generalized reward-seeking, we tested the effect of DO34 in a 2BC sucrose preference paradigm. DO34 had no effect on sucrose preference, suggesting a degree of specificity over natural rewards (**Fig. S6a**). Furthermore, given that depletion of 2-AG reduces EtOH drinking, we next tested if augmentation of 2-AG would increase EtOH drinking. The monoacylglycerol lipase inhibitor JZL-184 (10 mg/kg) was administered for three consecutive days to mice drinking 10% EtOH in the 2BC drinking paradigm. JZL-184 had no significant effect on EtOH preference or consumption (**Fig. S6b-c**) and did not affect body weight (**Fig. S6d**), indicating the effects of 2-AG modulation on EtOH drinking are not bidirectional.

### Pharmacological DAGL inhibition reduces aversion-resistant EtOH drinking

Alcohol seeking despite negative consequences is recognized as a key element of AUD and is a major obstacle to effective AUD treatment (2, 28). To determine whether DAGL inhibition could reduce EtOH consumption in a more clinically-relevant model, we utilized an aversion-resistant drinking model wherein animals actively consume EtOH that has been adulterated with the bitter tastant quinine (28). First, we validated this model by adulterating quinine in one of the two bottles after 4 weeks of the 2BC paradigm and determined the preference and consumption for the quinine adulterated bottles. Quinine was adulterated at 0.01, 0.03 and 0.1g/L in water, 10% EtOH and 20% EtOH bottles. Mice receiving water + quinine (0.03 and 0.1 g/L) exhibited very low preference (<7 %) (**Fig. S7a**), however, EtOH (10 and 20%) + quinine maintained a higher preference relative to the Water + quinine condition. Mice drinking 20% EtOH + quinine showed higher EtOH consumption compared to mice drinking 10% EtOH + quinine (**Fig. S7b**), leading us to use 20% EtOH for examining DO34 effects on aversion-resistant drinking. 0.03 g/L quinine was selected for further experiments, as mice exhibited robust aversion resistant drinking at this concentration (**Fig. S7a**).

To test DO34 effects, a separate cohort of 2BC mice was used and 0.03g/L quinine was added to either water bottles (control mice) or 20% EtOH bottles after 4 weeks of stable 2BC drinking. 20% EtOH + quinine showed higher preference compared to the water + quinine mice (Fig. 2a). DO34 (50 mg/kg) treatment significantly reduced the EtOH preference (**Fig. 2a**) and consumption (**Fig. 2b**). DO34 treatment had no effect on total fluid consumption (**Fig. S7c**). These data suggest DO34 can reduce aversion-resistant alcohol consumption.

**Figure 2:**
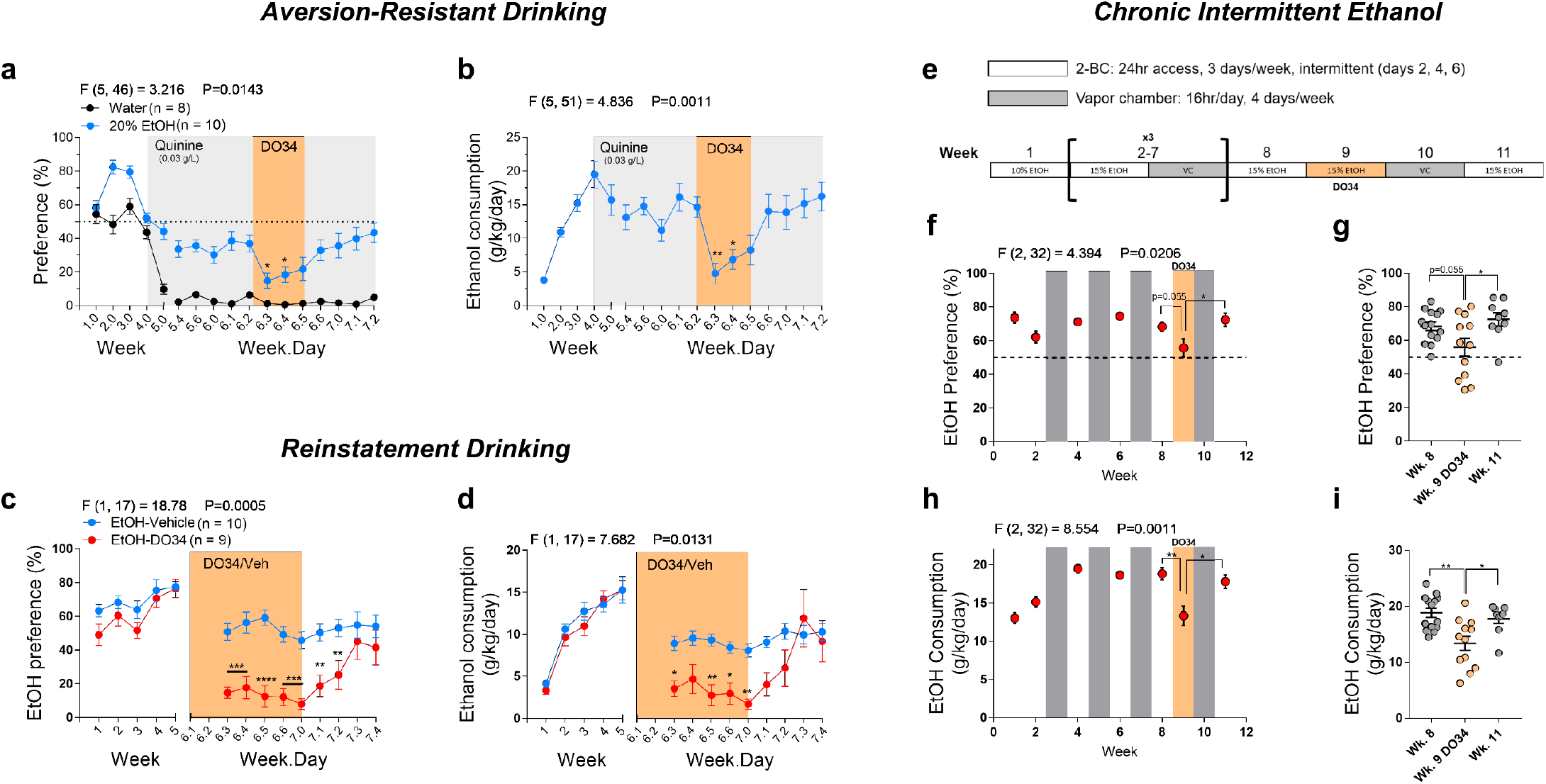
Pharmacological DAGL inhibition decreases EtOH intake across aversion-resistant, reinstatement, and chronic intermittent models of EtOH drinking. **(a)** DO34 treatment decreased preference and **(b)** consumption of EtOH adulterated with 0.03 g/L of the bitter tastant quinine (blue). Water + quinine control shown for reference (black) to demonstrate EtOH drinking persists despite aversive properties of quinine. **(c-d)** DO34 treatment reduced reinstatement of EtOH drinking after 5 weeks of drinking and 10 days of withdrawal. DO34-treated mice showed reduced **(c)** preference and **(d)** consumption upon re-initiation of EtOH drinking, relative to vehicle-treated mice. **(e)** Schematic depicting the chronic intermittent EtOH exposure paradigm. **(f)** DO34 treatment caused a non-significant reduction in EtOH preference (P=.055) in the chronic intermittent EtOH paradigm. **(g)** Graph depicting individual mouse EtOH preference during baseline, DO34 treatment, and recovery weeks. Preference during DO34 treatment was significantly lower compared to the recovery week. **(h)** DO34 treatment significant decreased EtOH consumption. **(i)** Graph depicting individual mouse EtOH consumption during baseline, DO34 treatment, and recovery weeks. EtOH consumption during DO34 treatment was significantly lower compared to both baseline and recovery weeks. EtOH data were analyzed by one-way ANOVA on time points 6.2 – 7.0 (to include baseline, drug treatment, and two recovery points) followed by a Holm-Sidak test for multiple comparisons to baseline control. **(c-d)** Data were analyzed by repeated measures two-way ANOVA followed by a Holm-Sidak test for multiple comparisons between treatment conditions. **(f-i)** Data analyzed by one-way ANOVA on time points 8, 9, and 11weeks followed by a Holm-Sidak test for multiple comparisons. *P* and *F* values for main drug effects of drug treatment reported on all graphs. Sample size *n* reported on graphs for experiments in **a-d**. *n* = 9-14 mice in **(f-i)**. Female mice were used for all experiments and all DO34 treatments are dosed at 50 mg/kg in **a-i**. Significance for post-hoc multiple comparisons reported on graphs (* P<.05, * * P<0.01, * * * P<0.001, * * * * P<0.0001). Data are mean ± SEM.

### Pharmacological DAGL inhibition reduces reinstatement of EtOH drinking

Another major obstacle to effective AUD treatment is the susceptibility to relapse after short or extended abstinence (29). We therefore used a model of EtOH reinstatement to test the effects of DO34 treatment on the potential for relapse after EtOH abstinence. Mice were subjected to our 2BC paradigm for 5 weeks (at 10% EtOH), followed by a 10-day withdrawal period where EtOH bottles were replaced with water. After the withdrawal period, mice were re-exposed to 10% EtOH for 9 days. Mice were treated with vehicle or DO34 (50 mg/kg) for 7 consecutive days, starting 2 days prior to the initiation of EtOH re-exposure. DO34-treated mice showed significantly lower EtOH preference (**Fig. 2c**) and consumption (**Fig. 2d**) compared to vehicle-treated mice upon EtOH re-exposure. DO34 treatment concomitantly increased total fluid consumption, again demonstrating the DO34-driven decrease in EtOH consumption is not due to off-target reductions in fluid intake observed with the control compound DO53 (**Fig. S7d**).

### Efficacy of DAGL inhibition is maintained following chronic intermittent EtOH exposure and dependence-like drinking

AUD exists on a severity spectrum and involves a shift from problematic alcohol use to dependence (2, 30). This transition is driven by a shift from positive to negative reinforcement (30), and thus may involve differing neurobiological mechanisms in early vs. severe, late-stage of AUD. Critical to the development of pharmacotherapies for AUD is the examination of relative efficacy during varying stages of AUD. To address this, we examined the efficacy of DO34 in mitigating alcohol consumption following a model of chronic intermittent EtOH (CIE) exposure to develop escalation and dependence-like drinking. CIE models are widely used to develop EtOH dependence in rodents and generally result in more severe AUD symptomology and higher intoxication levels relative to continuous access models (28, 31, 32). Mice were exposed to intermittent 2BC 15% EtOH drinking for 24hr periods, 3x per week. On alternating weeks, mice were exposed to EtOH vapor inhalation for 16 hr/day, 4x per week (**Fig. 2e**). Mice exhibited a clear escalation in EtOH preference and consumption following the first week of EtOH vapor exposure. Following 3 2BC-vapor chamber cycles and one additional week of baseline drinking, mice were treated daily with DO34 for 6 days, starting the day before the first drinking session. DO34 caused a small but non-significant (p=0.05) reduction in EtOH preference, and EtOH preference was significantly lower during DO34 treatment when compared to the recovery week after an additional 2BC-vapor cycle (**Fig. 2f-g**). DO34 treatment significantly reduced EtOH consumption during the treatment week and consumption recovered to baseline following an additional cycle of 2BC-vapor treatment (**Fig. 2h-i**). DO34 treatment also caused a small but significant reduction in total fluid consumption during the treatment week and after recovery (**Fig. S7e-f**). These data suggest that DAGL inhibition may be a broadly applicable therapeutic strategy for the treatment of varying stages of AUD, including late-stage dependence-like drinking modeled using the CIE exposure paradigm.

### Pharmacological DAGL inhibition decreases anxiety- and depressive-like behaviors during protracted abstinence

2 AG signaling is an important modulator of anxiety- and depressive-related behaviors (7– 9, 33, 34), and depletion of 2-AG produces anxiety-like phenotypes and exacerbates the affective consequences of stress exposure (9, 34–36). Additionally, 2-AG augmentation has been shown to mitigate anxiety-like behaviors in a rodent model of EtOH withdrawal (37). When considering these data in the context of DAGL inhibition for treating AUD, a potential problem with this therapeutic strategy is that pharmacological depletion of 2-AG may produce affective disturbances, similar to those previously observed with the CB_1_ receptor inverse agonist Rimonabant (20). We therefore wanted to examine the potential affective side effect profile of DAGL inhibition by examining the effects of DAGL inhibition on anxiety- and depressive-like behaviors during protracted abstinence. To test this, female mice were exposed to our 2BC paradigm at 20% EtOH (or water as a control) for six weeks and placed into forced abstinence for two weeks (**Fig. 3a-c**). Mice were then treated with vehicle or DO34 (50 mg/kg) on testing days two hours prior to each behavioral test (**Fig. 3a**). During this treatment period, mice were tested in the novelty-induced feeding suppression (NIFS), marble burying test, forced swim test (FST), and elevated plus maze (EPM) assays. EtOH abstinent animals treated with vehicle exhibited a higher latency to first bite relative to water control mice in the NIFS assay, suggesting an anxious phenotype produced by withdrawal from EtOH. DO34 treatment reduced this latency in EtOH abstinent but not water control mice, revealed by an EtOH treatment X DO34 interaction (**Fig. 3d**). In the marble burying assay, DO34 treatment reduced the number of buried marbles, regardless of EtOH history (**Fig. 3e**). Similarly, DO34 reduced immobility time in the FST in both water and EtOH mice (**Fig. 3f**). In the EPM assay, DO34 treatment increased open arm entries and decreased immobility time only in EtOH abstinent mice, again supported by an EtOH treatment X DO34 interaction (**Fig. 3g**). These data suggest that DO34 does not produce an anxiety- or depressive-like phenotype during protracted alcohol withdrawal and, in contrast, may exhibit anxiolytic or antidepressant properties, but only during alcohol withdrawal.

**Figure 3:**
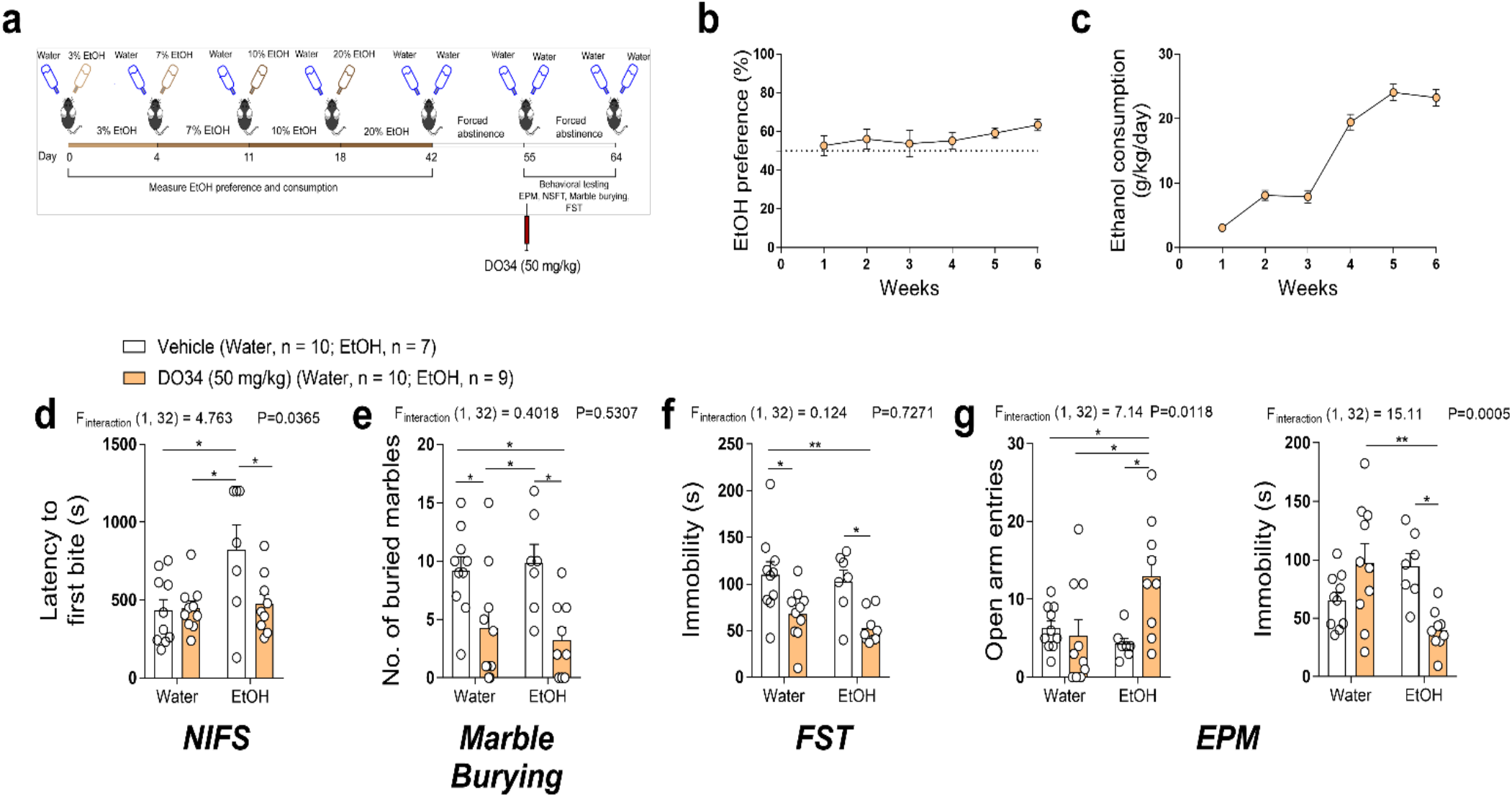
DO34 treatment does not cause negative affective phenotypes during protracted abstinence from EtOH drinking. **(a)** Schematic depicting 2BC drinking paradigm and time course of forced abstinence and behavioral testing. **(b)** EtOH preference and **(c)** consumption in mice prior to behavioral experiments. **(d)** EtOH drinking mice treated with vehicle showed higher latency to first bite in novelty-induced feeding suppression test and latency was reduced back to baseline in DO34-treated EtOH mice. **(e)** DO34 treatment decreased the number of buried marbles in water control and EtOH drinking mice. **(f)** DO34 treatment decreased immobility time in the forced swim test in water control and EtOH drinking mice. **(g)** DO34 treatment increased number of open arm entries and decreased immobility time in the elevated plus maze selectively in EtOH drinking mice. **(d-g)** Data were analyzed by two-way ANOVA followed by a Holm-Sidak test for multiple comparisons between all groups. Female mice were used in these experiments. *P* and *F* values for EtOH x drug interaction reported on graphs. Significance for post-hoc multiple comparisons reported on graphs (* P<.05, * * P<0.01). Data are mean ± SEM.

### Pharmacological DAGL inhibition decreases anxiety- and depressive-like behaviors during late chronic drinking

Given that AUD treatments may be initiated before a protracted abstinence period, a separate cohort of mice were subjected to the 2BC paradigm at 10% EtOH (or water control) and behavioral tests were performed at the end of 6 weeks of stable EtOH drinking, without an abstinence period (**Fig. 4a**). As done previously, mice were treated with vehicle or DO34 (50 mg/kg) 2 hours prior to each behavioral test. Mice were tested in the open field (OF), elevated zero maze (EZM), tail suspension test (TST), light-dark box, and 3-chamber social interaction assays. DO34 treatment decreased OF % center distance in water control mice, but not in EtOH mice (**Fig. 4b**). Additionally, DO34 treatment increased EZM open arm entries in EtOH mice, but not in water control mice (**Fig. 4c**). Total distance in the EZM was increased by DO34 treatment in both water and EtOH mice (**Fig. 4c**). In the TST, DO34 increased immobility time in water control mice, but decreased immobility time in EtOH mice, supported by a strong EtOH treatment X DO34 interaction (**Fig. 4d**). There was no effect of DO34 treatment or EtOH drinking on anxiety-like behaviors in the light-dark box or impairment in social function as measured by the 3-chamber social interaction test (**Fig. 4e-f**). These data are in agreement with the data collected from the protracted abstinence cohort. While some mild adverse phenotypes were observed in naïve mice, these data collectively suggest no adverse affective side effect liability from DAGL inhibition to reduce EtOH drinking after a history of EtOH consumption or abstinence in these mouse models studies.

**Figure 4:**
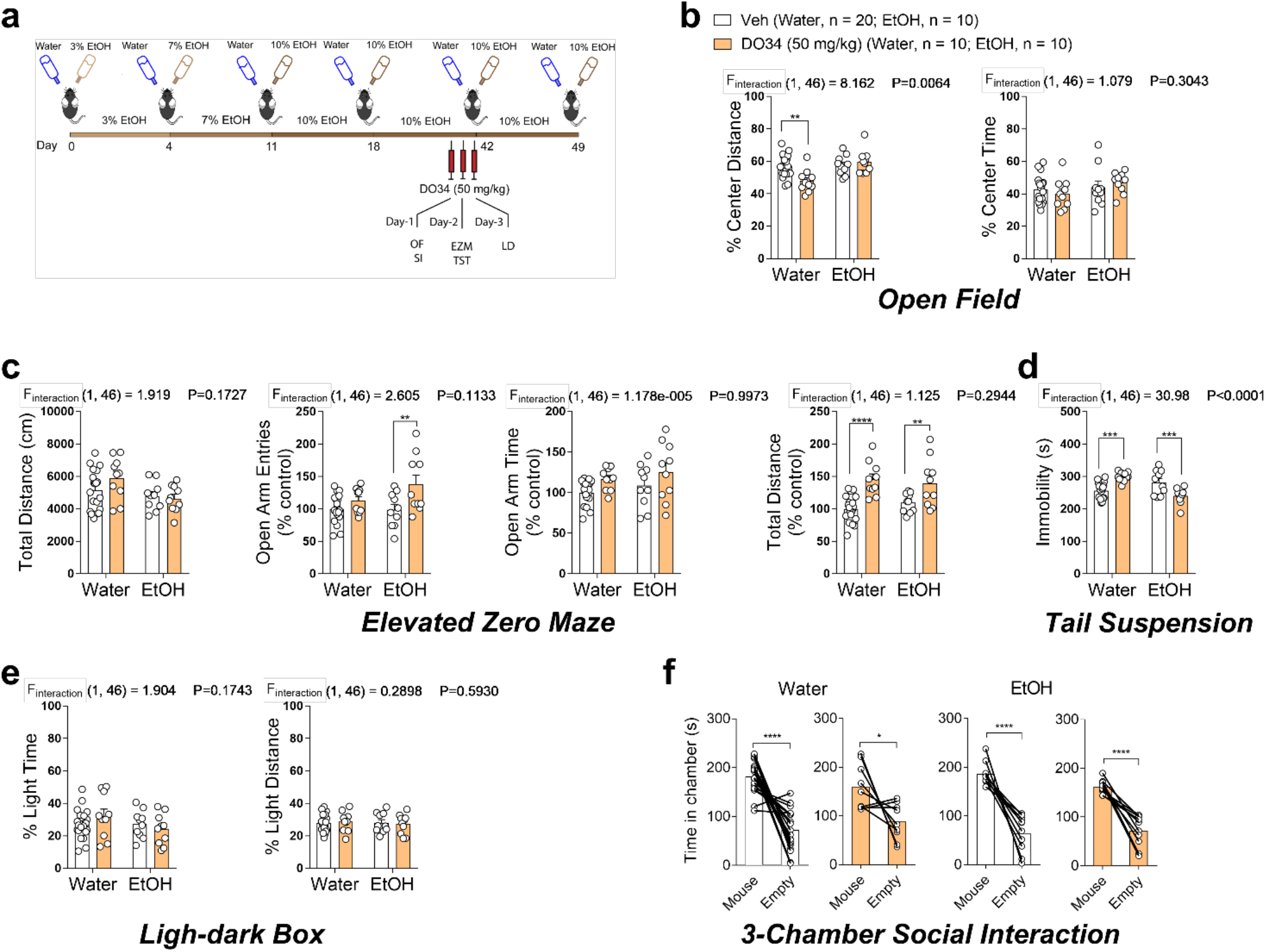
DO34 treatment does not cause negative affective phenotypes during late chronic EtOH drinking. **(a)** Schematic depicting 2BC drinking paradigm and time course of behavioral testing. **(b)** DO34 treatment decreased open field test % center time in water control mice but not in EtOH drinking mice. **(c)** DO34 treatment increased % open arm entries in EtOH mice but not water control mice and increased total distance travelled in both water and EtOH dunking mice. **(d)** DO34 increased immobility time in the tail suspension test in water control mice but decreased immobility time in EtOH mice. **(e)** DO34 treatment had no effect on anxiety-like behaviors in the light-dark box assay. **(f)** DO34 treatment had no effect on social behaviors in the 3-chamber social interaction test. **(b-e)** Data were analyzed by two-way ANOVA or **(f)** paired t-test followed by a Holm-Sidak test for multiple comparisons between all groups. Female mice were used in all experiments. *P* and *F* values for EtOH x drug interaction reported on graphs. Significance for post-hoc multiple comparisons reported on graphs (* P<.05, * * P<0.01, * * * P<0.001, * * * * P<0.0001). Data are mean ± SEM.

## DISCUSSION

Here we show that genetic and pharmacological inhibition of DAGL activity reduced alcohol consumption and preference across a range of distinct drinking models. Specifically, pharmacological DAGL inhibition and DAGLα KO mice show reduced voluntary alcohol consumption and preference under a continuous access model. Importantly, pharmacological DAGL inhibition also reduced alcohol consumption and preference in an aversion-resistant drinking model, a model of reinstatement drinking, and in a dependence-like model that leverages chronic intermittent vapor exposure to drive an escalation in voluntary alcohol consumption. In line with our data, DAGL inhibition by DO34 has been previously shown to reduce operant responding for alcohol (38). The magnitude of effect on EtOH intake observed after DAGL inhibition is comparable to or greater than those observed with clinically available treatments such as Naltrexone (39– 41) and Acomprasate (41, 42). Moreover, pharmacological DAGL inhibition did not affect sucrose preference, suggesting a lack of generalized effect on natural reward, although female DAGLα KO mice do exhibit reductions in sucrose preference under some conditions (35). Although DO34 inhibits both α and β isoforms of DAGL (22), the ability of DAGLα genetic knockout to reduce alcohol consumption suggests the α isoform may be the relevant molecular target contributing to the efficacy of DO34. Furthermore, we found a control compound DO53, which does not inhibit DAGL α or β, but does inhibit a number of off-targets also inhibited by DO34 (22), produced a transient suppression of alcohol intake that was paralleled by a reduction in non-specific fluid intake. This effect may be due to activity at shared off-targets or unique off-targets of DO53 such as ABHD3 (22). Regardless of the precise mechanisms by which DO53 transiently suppresses non-specific fluid intake behavior, it did not affect alcohol preference or alcohol consumption after repeated injection, providing an additional degree of confidence that the effects of DO34 on alcohol consumption and preference were mediated via DAGL inhibition. Overall, these data suggest targeting DAGLα may represent a novel approach to the treatment of AUD across a spectrum of severity and development of potent and selective DAGLα inhibitors should remain a high priority. In this context, that DAGLα transcript is known to be preferentially expressed within the human brain, while DAGLβ is more ubiquitously expressed (see **Fig. S8**), suggests selectively targeting DAGLα may have less somatic adverse effect liability, however this remains to be determined experimentally.

eCB signaling has been shown to regulate VTA dopamine neuron activity via suppression of afferent GABAergic drive (24–27). For example, 2-AG mediated suppression of GABA transmission onto VTA DA neurons is weaker in alcohol Sardinian alcohol-preferring rats relative to non-preferring rats (25, 27), an effect mediated by enhanced 2-AG degradation (25). These data suggest that 2-AG can promote alcohol preference via activity-dependent suppression of GABAergic transmission onto VTA dopamine neurons. Based on these data we tested the hypothesis that pharmacological DAGL inhibition could affect alcohol modulation of GABAergic transmission onto VTA DA neurons, which have bene shown to increase their firing rate in response to exogenous alcohol (16). Consistent with these data, *ex vivo* alcohol application to VTA brain slices decreased IPSC frequency into VTA dopamine neurons, as shown previously (23). This effect was blocked by DO34, suggesting DAGL inhibition could reduce alcohol consumption and preference via reductions in alcohol-stimulated dopamine neuron activity via maintaining high relative levels of GABAergic inhibition. Similar mechanisms have been proposed to underlie the effects of DAGL signaling on nicotine activation of VTA dopamine neurons (24). While these data suggest DAGL inhibition could reduce positive reinforcement-driven alcohol consumption by reducing alcohol-induced activation of VTA dopamine neurons, negative reinforcement driven mechanisms are thought to mediate alcohol seeking and drinking under dependent conditions (30). Although DO34 was able to reduce alcohol consumption in a dependent-like drinking model, the underlying synaptic mechanisms subserving these effects may indeed be different. Future studies should be aimed at elucidating the distinct mechanisms by which DAGL inhibition affects positive vs. negative reinforcement-driven alcohol drinking.

Another key finding of the present work is that DO34 did not increase anxiety or depressive-like behaviors in mice exposed to chronic alcohol or during protracted abstinence. In chronically drinking mice, DO34 reduced immobility in the TST, while it increased immobility in water drinking controls. These data exemplify the notion that the effects of DO34 are dependent on alcohol history and could explain, in part, the apparently contradictory results relative to previously published work that would predict an increase in affective behaviors (9, 36). For example, DO34 increases anxiety-like behaviors in mice exposed to acute stress (36) and impairs extinction of conditioned fear memories (43). Moreover, DAGLα KO mice exhibit anxiety-like behaviors under some, but not all, conditions, and some reports are conflicting. For example, one report found a robust anxiety-like phenotype in DAGLα KO mice in the light-dark box assay (35), but this effect was absent in DAGLα mice in a later report using a similar light-dark box assay (44). Furthermore, DO34 exerted anxiolytic effects during protracted abstinence, which was unexpected given the anxiogenic effects of DO34 observed after stress exposure (36).

These data again highlight the notion that behavioral function of 2-AG signaling (and thus the effects of DO34) on anxiety and depressive-like phenotypes is highly dependent upon context, such as alcohol use history. The mechanisms by which DO34 exerts unexpected anxiolytic effects remain to be determined. Interestingly, we recently showed that augmenting 2-AG levels via inhibition of the 2-AG degrading enzyme monoacylglycerol lipase (MAGL) also showed anxiolytic effects (9, 36), and MAGL inhibition has exhibited anxiolytic effects in alcohol withdrawal as well (37). This apparently paradoxical finding has also been observed in other models; for example, both inhibition (43) and augmentation (45) of 2-AG signaling can facilitate acute fear responses in conditioned fear paradigms. These data collectively suggest that bidirectional manipulations of 2-AG levels can have overlapping effects in some cases, and the behavioral effects of pharmacological 2-AG modulation may be highly context-dependent. The distinct mechanisms underlying these unusual effects of 2-AG modulation remain to be determined. Furthermore, there are several genetics variants that strongly alter DAGLα expression in the brain (see **Fig. S9** for one example, rs11604261), demonstrating that the baseline expression of DAGLα can vary across individuals. This suggests that variation within the *DAGLA* gene may demonstrate correlations with susceptibility to AUD or serve as a biomarker for predicting responsiveness to pharmacological DAGL inhibition. Large scale genetic studies are needed to assess this connection.

In summary, our data present a previously undescribed role for DAGL signaling in voluntary alcohol drinking, and posit targeting 2-AG synthesis as a promising pharmacotherapeutic mechanism for the treatment of AUD. The finding that DO34 does not produce affective phenotypes (and exhibits mild anxiolytic effects in some cases) further highlights the therapeutic potential of DAGL as a target for AUD and is especially pertinent given the neuropsychiatric side effects of CB_1_ receptor blockade (20). The variety of models in which we tested the effects of DO34 address several key hallmarks of AUD that serve as barriers to current treatments, including aversion-resistant, compulsive-like drinking, susceptibility to relapse, and dependence-like drinking. The ability of DO34 to reduce drinking in all these contexts suggest DAGL inhibition may exhibit therapeutic utility for treating the broad spectrum of mild to late-stage, severe AUD, which could have broad implications. Future studies should be aimed at determining the cellular and synaptic mechanisms underlying the effects of DO34 on early vs. late, dependence-like drinking, as well as on the development of more selective DAGLα inhibitors to further test this hypothesis and ultimately lead to the discovery of more safe and effective therapies for AUD.

## METHODS

### Subjects

C57BL/6j male and female mice between 6–8 weeks of age at the beginning of experiments were used. DAGLα^−/−^ male and female mice were generated by disruption of exon 8 and were maintained on a C57Bl6/N background by interbreeding DAGLα+/− mice. Further details are described in Ref. (35). Tyrosine hydroxylase (TH) reporter mice (TH::Ai14 mice) were generated by crossing TH-Cre mice (Jax Stock No: 008601) with Ai14 mice (Jax Stock No: 007914) for Cre-dependent Td-tomato expression in TH+ cells to label catecholaminergic neurons. All mice were group housed on a 12:12 light-dark cycle (lights on at 6:00 a.m.) with food and water available ad libitum. All behavioral testing was performed between 6:00 am and 6:00 pm.

### Drugs and treatment

The DAGL inhibitor DO34 (50 mg/kg; Glixx Laboratories Inc., MA, USA), control compound DO53 (50 mg/kg; synthesized in-house, see **Fig. S1**), MAGL inhibitor JZL184 (10 mg/kg; Cayman Chemical, MI, USA), pyrazole (68 mg/kg; Sigma-Aldrich, WI, USA) and quinine hydrochloride (0.01, 0.03 and 0.1 g/L; Millipore, MA, USA) were used. DO34 and DO53 (or vehicle control) were administered by intraperitoneal (i.p.) injection at a volume of 10 ml/kg in a formulation containing ethanol (Pharmco, KY, USA): kolliphor (Sigma-Aldrich, WI,): saline (Hospira, IL, USA) (1:1:18). JZL184 (or vehicle control) was administered by i.p. injection at a volume of 1 ml/kg in DMSO (Sigma-Aldrich, WI, USA, Cat. No. D8414). EtOH + pyrazole solutions for vapor chamber experiments were prepared in saline and injected i.p. at a dose of 1.6 g/kg EtOH + 68 mg/kg pyrazole. Drug pretreatment times were two hours prior to behavioral testing or 30 minutes prior to vapor chamber sessions. Drugs for EtOH drinking experiments were injected 2 hours before starting the dark cycle. Quinine hydrochloride (Millipore, MA, USA) was added to the drinking water or ethanol bottles at varying concentrations. 190 proof ACS/USP grade grain-derived EtOH was used to make EtOH solutions.

### Two-bottle choice (2BC) EtOH drinking paradigm

Mice were singly housed and acclimatized for 5-7 days in 2BC cages. Mice had access to two sippers throughout the experiment. For EtOH drinking mice, EtOH concentration was slowly increased from 3% to 10% or 20% as shown in **Figure 1a** and maintained on respective solutions for the duration of the experiment. Water and ethanol intake were monitored either daily, after 4 days, or weekly depending on the experimental timeline. For the intermittent access model (see **Fig. 2e**), mice were given intermittent access to alcohol (10% on week one, 15% thereafter) 3 days per week for 24-hour periods on alternating days. Whenever mice were given pharmacological treatments, water intake, EtOH intake and body weights were measured daily. Food was provided *ad libitum* throughout the alcohol drinking paradigm. EtOH naive mice (control) were also housed in the same conditions, but two bottles of water remained in their cage until day of experiment. EtOH preference was determined as ((EtOH intake (g)/total fluid intake (g)) × 100). EtOH consumption was determined as [((EtOH intake (g)/mice body weight (kg)/number of days) × EtOH concentration × EtOH density (0.816 g/mL)]. Dummy cages were used to calculate water and EtOH drip and subtracted from intake values.

### Aversion-Resistant EtOH drinking model

After 4 weeks of EtOH drinking in 2BC EtOH drinking paradigm, quinine was added to either water (control mice) or EtOH bottles (EtOH group) at varying concentration (0.01, 0.03 and 0.1 g/L). EtOH quinine and water quinine preference and consumptions were calculated as described above.

### Ethanol vapor inhalation treatment for chronic intermittent EtOH (CIE) exposure

For chronic intermittent ethanol exposure, female C57Bl6/J were exposed to vapor inhalation for 16 hours per day, 4 days per week, on alternating weeks as shown in **Fig. 2e**. Mice were given i.p. injections of EtOH (1.6 g/kg) + pyrazole (alcohol dehydrogenase inhibitor, 68 mg/kg) 30 minutes prior to the start of each session. Mice were then placed in a chamber with volatilized EtOH (18-22 mg/L) for 16 hours. Mice had *ad libitum* access to food and water for the duration of the session. These procedures were similar as done previously (46) which were able to produce blood EtOH levels of 150–185 mg/dl. Following 4 days of vapor treatment, mice went through 72 hours of abstinence before returning to the intermittent 2BC paradigm described above.

### EtOH Reinstatement Drinking

After 5 weeks of EtOH drinking in 2BC EtOH drinking paradigm, ETOH bottles were replaced with water bottles for 10 days. After 10 days, EtOH was provided to study reinstatement of EtOH drinking behavior. EtOH preference and consumptions were calculated as described above.

### Behavioral experiments

#### Novelty-induced feeding suppression (NIFS)

NIFS assay was performed as described previously (33). Briefly, mice were deprived of food for a 48hr period, with brief access to food from 23-25hr. Following food deprivation, mice were acclimated to the testing room for at least 2hr prior to placement in a brightly lit (250-300 lux) 50 x 50cm arena with a single food pellet in the center of the arena. Mouse movement and behavior was tracked with an overhead camera using AnyMaze software (Stoelting, IL, USA). Mice remained in the arena until their first bite of the food pellet. Mice were given *ad libitum* access to food after testing.

#### Marble burying test

For the marble burying test, empty cages without food or water were filled with 5cm fresh Diamond Fresh Soft Bedding (Envigo, IN, USA) and 20 marbles (5 rows, 4 columns) placed on top of the bedding. Mice were placed in the cage for 30 minutes, then removed. Marbles were considered “buried” if 2/3 of the marble was covered by bedding. Marbles were counted manually by an experimenter that was blinded to the treatment condition. Light intensity in the room was 200-250 lux.

#### Forced swim test (FST)

FST was performed as described previously (33). Briefly, mice acclimated to the testing room for at least 2 hours then were placed in a cylinder containing water (23-25°C) at a level such that mice could not touch the bottom or the top. The test lasted for 6 minutes and filmed via overhead camera and later scored by an observer blinded to treatment. Immobility time was analyzed during the last 4 minutes of the test.

#### Elevated plus maze (EPM)

EPM assay was performed similar to as described previously (34). The apparatus consisted of two open arms (30 × 10cm) and two closed arms (30 × 10 × 20cm) that met at a center junction (5 × 5cm) and was elevated 50cm above the floor. Light intensity in the open arms was 200-250 lux and <100 lux in the closed arms. Mice were placed in the center of the apparatus, facing an open arm and allowed to freely explore for 6 minutes. Movement and behavior were tracked via an overhead camera and AnyMaze software (Stoelting, IL, USA).

#### Open field

For open-field testing (OFT), exploration of a novel open field arena contained within a sound-attenuating chamber was monitored for 30 min (27.9 × 27.9 × 20.3 cm; MED-OFA-510; MED Associates, St. Albans, Vermont). The walls of the open field arena were made of clear plexiglass; this arena was contained within an opaque sound-attenuating chamber. Beam breaks from 16 infrared beams were recorded by Activity Monitor v5.10 (MED Associates) to monitor position and behavior.

#### 3-chamber social interaction

Social behavior was tested in a 3-chamber acrylic arena with opaque outer walls and clear walls separating each 20 cm x 40 cm chamber, which contained 10 cm sliding doors for introduction of the test mouse into the middle chamber. Testing was performed under low lighting conditions (∼40 lux) and mice were acclimated to this lighting for at least 1 hour before testing began. The test mouse was allowed to explore the apparatus, which contained one empty inverted wire pencil cup in each outer chamber, for 10 minutes and was then coaxed back into the middle chamber and doors replaced. The target mouse (a female WT C57Bl6 mouse, which was aged matched and previously acclimated to the pencil cup for 30 min) was then placed under one of the pencil cups and the doors were removed allowing the test mouse to explore the entire apparatus for 5 minutes. The mouse was monitored by an overhead camera and AnyMaze software (Stoelting, IL, USA) and the time spent in each chamber was measured. Preference for the social target was determined within each group by comparing the time spent in the target-mouse chamber to the time spent in the empty pencil cup chamber using a paired two-tailed student t-test. Mice that did not spend significantly more time in the target-mouse chamber were considered deficient in sociability.

#### Elevated-zero maze

The elevated-zero maze (EZM, San Diego instruments, California, USA) is annular white platform and divided four equal quadrants. It consisted of two open arms and two closed arms. The outer and inner diameter of EZM was 60.9 cm and 50.8 cm, respectively. The apparatus was elevated 60.9 cm from the floor. Light levels in the open arms were approximately 200 lux, while the closed arms were <100 lux. Mice were placed in the closed arm of the maze and allowed to explore for 5 min. ANY-maze (Stoelting, Wood Dale, Illinois, USA) video-tracking software was used to monitor and analyze behavior during the test.

#### Tail suspension test

Each mouse was suspended from its tail by using adhesive tape on a flat metal plate for 6 min. In order to prevent animals observing or interacting with each other, each mouse was suspended within its four-walled rectangular compartment (33 x 31.75 x 33 cm). The mouse is suspended in the middle of the compartment and the width and depth are sufficiently sized so that the mouse cannot make contact with the walls. Immobility time was automatically measured by the Med Associates Inc. tail suspension software.

#### Light-dark box test

The light-dark test was performed as previously described (9). Mice were individually placed into sound-attenuating chambers (27.9 × 27.9 cm; MED-OFA-510; MED Associates, St. Albans, VT, USA) containing dark box inserts that split the chamber into light (250-400 lux) and dark (<5 lux) halves (Med Associates ENV-511). Beam breaks from 16 infrared beams were recorded by Activity Monitor v5.10 (MED Associates) to monitor position and behavior during the 10-min testing period.

### Electrophysiology Experiments

Male and female TH::Ai14 mice (described above) were used for electrophysiological experiments. Acute slice preparation was performed as described previously (36). Coronal midbrain slices were cut at 200µm. Following recovery, slices were transferred to a recording chamber and perfused at a rate of 2-3mL/minute with oxygenated artificial cerebrospinal fluid (ACSF; 31-33°C) recording solution consisting of (in mM): 113 NaCl, 2.5 KCl, 1.2 MgSO_4_·7H_2_0, 2.5 CaCl_2_·6H_2_0, 1 NaH_2_PO_4_, 26 NaHCO_3_, 20 glucose, 3 Na-pyruvate, and 1 Na-ascorbate. TH+ cells were visually identified by their red fluorescence. Whole-cell recordings of spontaneous inhibitory post-synaptic currents (sIPSCs) were performed in voltage clamp configuration at −70mV in the presence of 10µM CNQX and 30µM D-AP5. 3–6 MΩ borosilicate glass pipettes were filled with the intracellular pipette solution consisting of (in mM): 125 KCl, 4 NaCl, 10 HEPES, 4 MgATP, 0.3 NaGTP, 10 Na-phosphocreatine, and 0.2 EGTA. Following break in, all cells were allowed to dialyze with the pipette solution for 3 minutes prior to recordings. Cells with an access resistance of ≤25 MΩ were included in analyses. DO34 was dissolved in DMSO stocks at 10mM and diluted in ACSF for a final concentration of 2.5µM. DO34-treated slices were incubated for a minimum of 45 minutes prior to recordings. sIPSC baselines were recorded for 2 minutes, followed by 10-minute bath application of 100 mM ethanol. Cells were then recorded for an additional 2 minutes in the continued presence of EtOH. Recordings were performed using a MultiClamp 700B amplifier (Molecular Devices, CA, USA), and Clampex software (Molecular Devices, CA, USA). Traces were analyzed using pCLAMP 10 software (Molecular Devices, CA, USA).

### Statistics

Statistical significance was calculated using a one-way or two-way ANOVA with post hoc Holm-Sidak’s multiple comparisons test as noted in figure legends. All statistical analyses were conducted using GraphPad Prism 7 (GraphPad Sofrware, CA, USA). For behavioral studies, all replicates (n values) represent biological replicates defined as data derived from a single mouse. Data are presented as mean ± S.E.M. unless otherwise stated in the figure legends. Significance was determined by p<0.05 throughout the manuscript. F and P values for ANOVA analyses are indicated within figure panels, while post hoc significance level is indicated above individual bars or time points. Testing was counterbalanced, but no randomization was performed, and sample sizes were derived empirically during the course of the experiments and guided by our previous work using these assays.

### Study Approval

All studies were carried out in accordance with the National Institutes of Health Guide for the Care and Use of Laboratory Animals and approved by the Vanderbilt University Institutional Animal Care and Use Committee (M1600213-01; M1800046).

## AUTHOR CONTRIBUTIONS

G.B., and N.D.W., and S.P. conceived the study, designed experiments, and co-wrote the manuscript.; S.W.C and D.G.W. assisted in alcohol drinking model development and optimization.; G.B., N.D.W., T.A.P., A.A., and V.R.M. performed experiments and acquired and analyzed data in the laboratories of S.P. and D.J.W.; J.U. synthesized the control compound DO53 in the laboratory of L.J.M.; D.C.S. acquired gene expression data.

## ACKNOWLEDGEMENTS

These studies were supported by NIH grants AA026186 (S.P.) and AA019455 (D.G.W.) and Brain Behavior Research Foundation NARSAD Young Investigator Awards 27172 (S.W.C.).

