## Supplemental Materials for "Targeting Diacylglycerol Lipase to Reduce Alcohol Consumption"

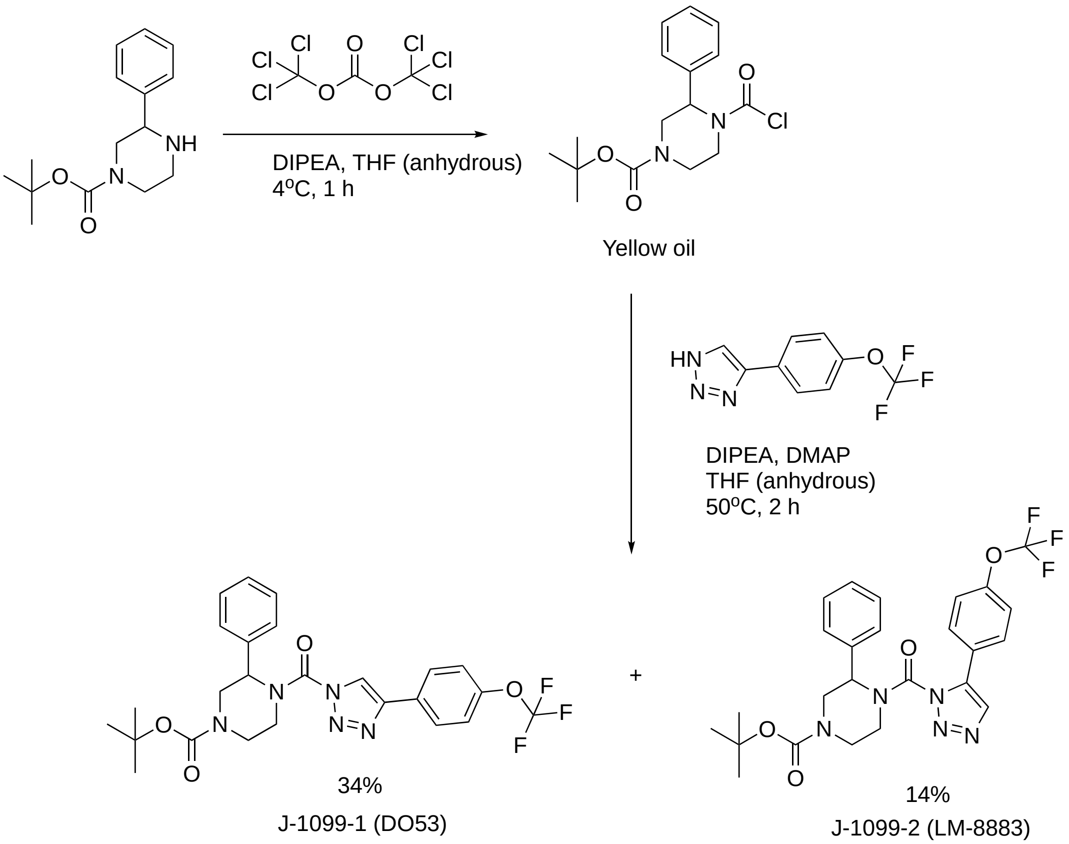


**Figure S1: Chemical synthesis and spectroscopic characterization of (4-{[(2-Methyl-2-propanyl)oxy]carbonyl}-2-phenylpyperazinyl){4-[(4- trifluoromethoxy)phenyl]-1H-1,2,3-triazol-1-yl}metanone (DO53).**

*Synthetic Procedure*: Diisopropylethylamine (275 mg, 2 mmol) followed by triphosgene (100 mg, 0.34 mmol) was added to a solution of 1-Boc-3-phenylpiperazine (175 mg, 0.65 mmol) dissolved in anhydrous tetrahydrofuran (5 mL), and the resultant reaction mixture was stirred 1 hour on ice. The reaction mixture was poured into water (5 mL) and extracted with ethyl acetate, and the organic layer was washed with water and brine, dried over anhydrous sodium sulfate and concentrated in vacuo. The residue was kept under vacuum overnight, then dissolved in anhydrous tetrahydrofuran (10 mL), to which diisopropylethylamine (275 mg, 2 mmol), 4-(dimethylamino)pyridine (80 mg, 0.65 mmol) and 4-(4- trifluoromethoxyphenyl)-1H-1,2,3-triazole (150 mg, 0.65 mmol) were added chronologically. The reaction mixture was wormed and stirred for 2 hours at 50 °C. The reaction mixture was cooled to ambient temperature and poured into saturated aqueous ammonium chloride solution and extracted with ethyl acetate. Combined organic layer was washed with water and brine, dried over anhydrous sodium sulfate and concentrated in vacuo. The crude product was purified by silica gel column chromatography (ethyl acetate : hexane = 1:8, v/v) to afford (4-{[(2-methyl-2-propanyl)oxy]carbonyl}-2-phenylpyperazinyl){4-[(4- trifluoromethoxy)phenyl]-1H-1,2,3-triazol-1-yl}metanone (DO53) (119 mg, 34%).

*Spectroscopic Characterization*: $^{1}$H NMR (DMSO-$d_6$ 600 MHz): 1.33 (s, 9H), 3.03-3.20 (m, 1H), 3.21-3.42 (m, 1H), 3.45-3.59 (m, 1H), 3.69-3.89 (m, 1H), 3.95-4.10 (m, 1H), 4.45-4.55 (m, 1H), 5.58 (s, 1H), 7.25−7.35 (m, 1H), 7.36−7.42 (m, 4H), 7.48 (d, 2H, $J$ = 8.6 Hz), 8.08 (d, 2H, $J$ = 8.6 Hz), 8.19 (s, 1H). ESI MS calcd. $C_{25}H_{26}F_3N_5O_4$ $[M+H]^+$ 518.20, MS found 518.08.

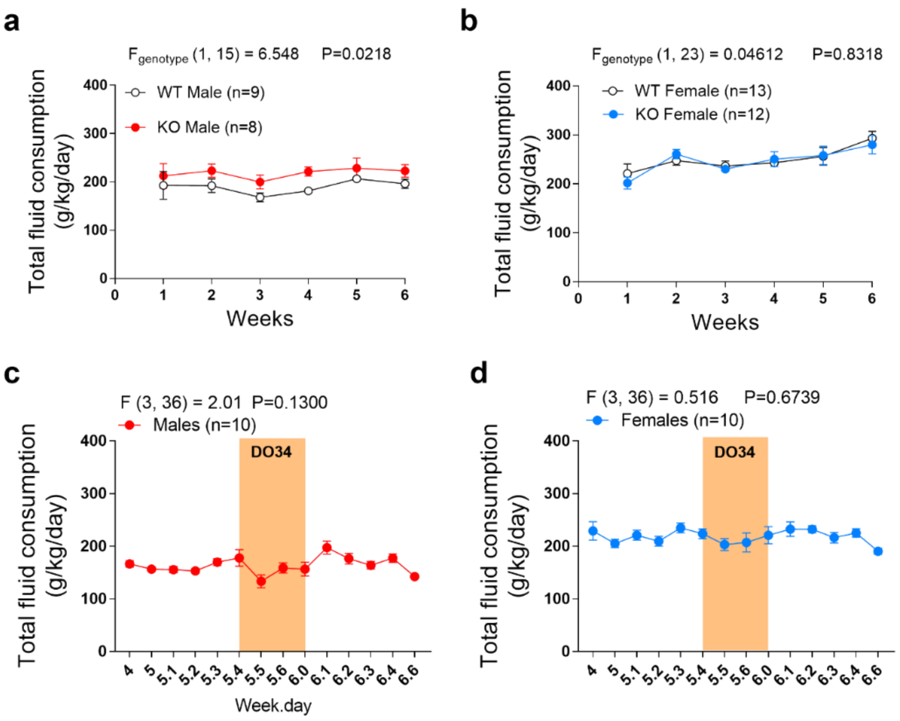


**Figure S2: DAGL inhibition does not decrease fluid consumption.** Genetic deletion of DAGLα does not decrease total fluid consumption in **(a)** male or **(b)** female DAGLα$^{-/-}$ mice. DO34 treatment does not decrease total fluid consumption in **(c)** male or **(b)** female mice. All DO34 treatments were dosed at 50 mg/kg. **(a-b)** Data analyzed by repeated measures two-way ANOVA followed by a Holm-Sidak test for multiple comparisons between genotypes or **(c-d)** one-way ANOVA on time points 5.4 – 6.2 (to include baseline, drug treatment, and one recovery point) followed by a Holm-Sidak test for multiple comparisons to baseline control. Sample size *n*, *P*, and *F* values for main effects of genotype or drug treatment reported on graphs. Significance for post-hoc multiple comparisons reported on graphs (*P<.05, **P<0.01, ***P<0.001, ****P<0.0001). Data are mean ± SEM.

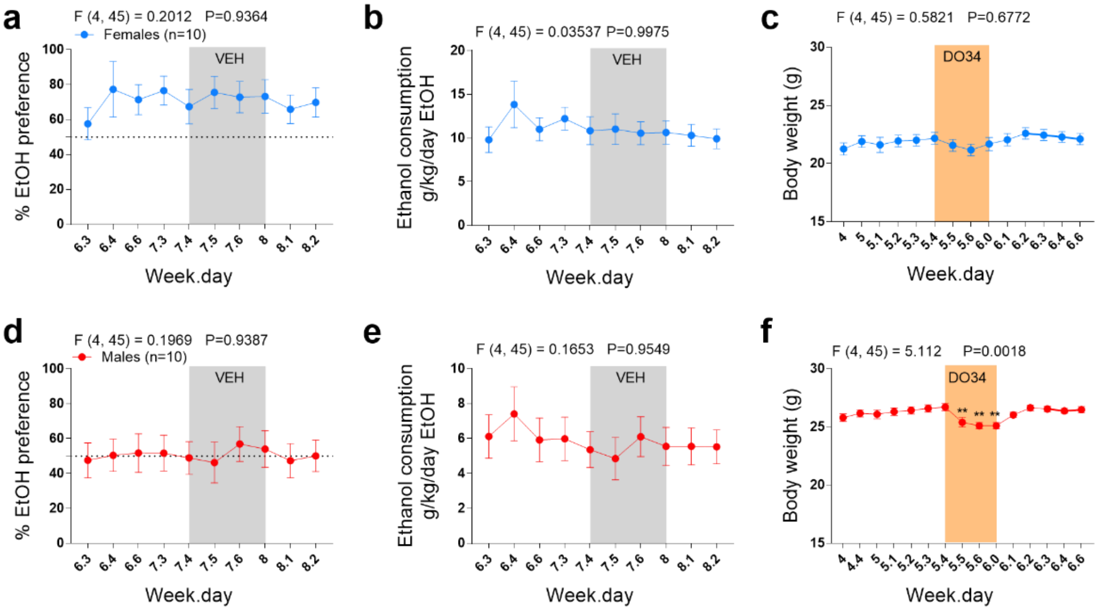


**Figure S3: Vehicle treatment does not affect EtOH preference or consumption and DO34 reduces body weight only in male mice.** Vehicle treatment had no effect on EtOH **(a,d)** preference or **(b,e)** consumption in either sex. **(c)** DO34 had no effect on body weight in female mice. **(f)** DO34 decreased body weight in male mice. Body weight recovered after cessation of DO34 treatment. All DO34 treatments were dosed at 50 mg/kg. Data were analyzed by one-way ANOVA on time points 7.4 – 8.1 (to include baseline, drug treatment, and one recovery point) followed by a Holm-Sidak test for multiple comparisons to baseline control. Sample size *n*, *P*, and *F* values for main effects of drug treatment reported on graphs. Significance for post-hoc multiple comparisons reported on graphs (*P<.05, **P<0.01, ***P<0.001, ****P<0.0001). Data are mean ± SEM.

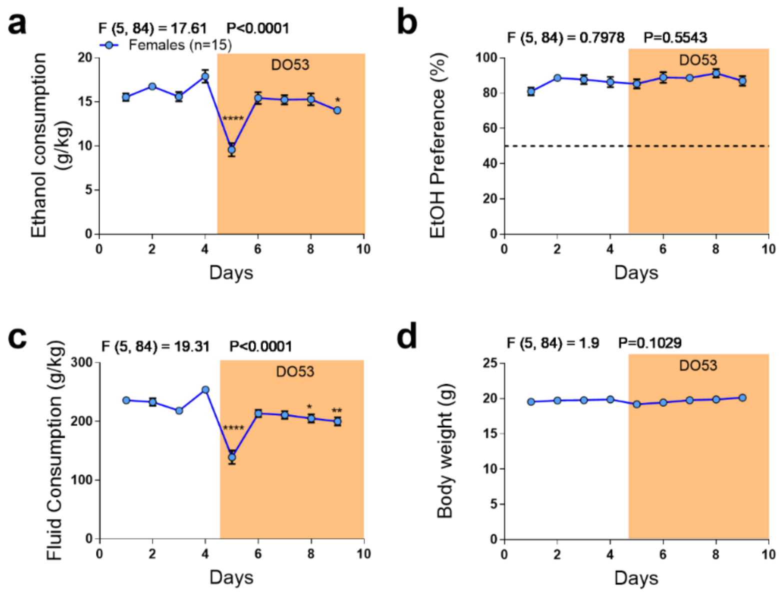


**Figure S4: DO53 treatment concomitantly reduces EtOH and total fluid consumption with no effect on EtOH preference on body weight. (a)** Treatment with the control compound DO53 caused a transient reduction in EtOH consumption. (b) DO53 had no effect on EtOH preference. (c) DO53 treatment caused a transient reduction in total fluid consumption, paralleling the effect on EtOH consumption. (d) DO53 treatment had no effect on body weight. Baseline data were averaged for analyses. Data were analyzed by one-way ANOVA on averaged baseline and individual treatment days followed by a Holm-Sidak test for multiple comparisons to averaged baseline control. Sample size *n*, *P*, and *F* values for main effects of drug treatment reported on graphs. All DO53 treatments were dosed at 50 mg/kg. Female mice were used for these experiments. Significance for post-hoc multiple comparisons reported on graphs (*P<.05, **P<0.01, ***P<0.001, ****P<0.0001). Data are mean ± SEM.

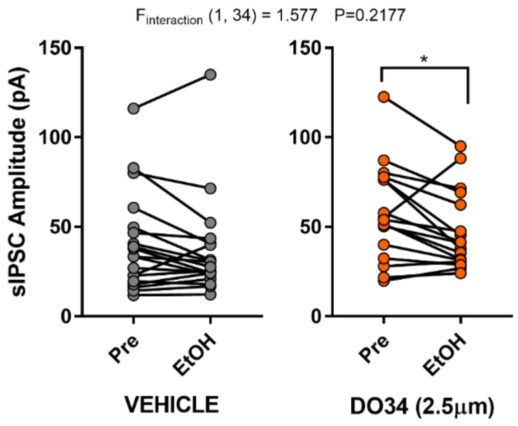


**Figure S5: EtOH reduces sIPSC amplitude onto putative dopamine neurons in the posterior VTA of DO34-treated slices.** Bath application of 100mM EtOH reduced sIPSC amplitude in DO34- but not vehicle-treated slices. All cells recorded were putative dopamine neurons visually-identified by their red fluorescence (see main text, **Fig. 1k**). Data were analyzed by repeated measures two-way followed by a Holm-Sidak test for multiple comparisons between baseline and EtOH treatment. Vehicle, $n$ = 19 cells; DO34, $n$ = 17 cells; from 13 mice. $P$ and $F$ value for EtOH x DO34 interaction reported on graph. Significance for post-hoc multiple comparisons reported on graph (*P<.05).

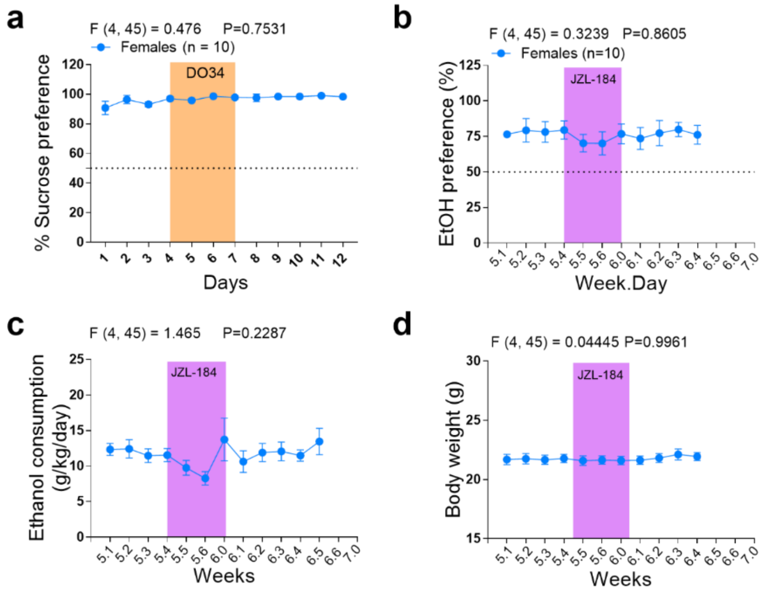


**Figure S6: DO34 does not alter sucrose preference and 2-AG augmentation has no effect on EtOH drinking or body weight. (a)** DO34 treatment had no effect on 2BC sucrose preference. (b) Treatment with the monoacylglycerol lipase inhibitor JZL-185 had no effect on EtOH preference, (c) EtOH consumption, or (d) body weight. Data were analyzed by one-way ANOVA on time points 4 – 8 (a) or 5.4 – 6.1 (to include baseline, drug treatment, and one recovery point) followed by a Holm-Sidak test for multiple comparisons to baseline control. Sample size *n*, *P*, and *F* values for main effects of drug treatment reported on graphs. All DO34 treatments were dosed at 50 mg/kg. All JZL-184 treatments were dosed at 10 mg/kg. Female mice were used for these experiments. Significance for post-hoc multiple comparisons reported on graphs (*P<.05, **P<0.01, ***P<0.001, ****P<0.0001). Data are mean ± SEM.

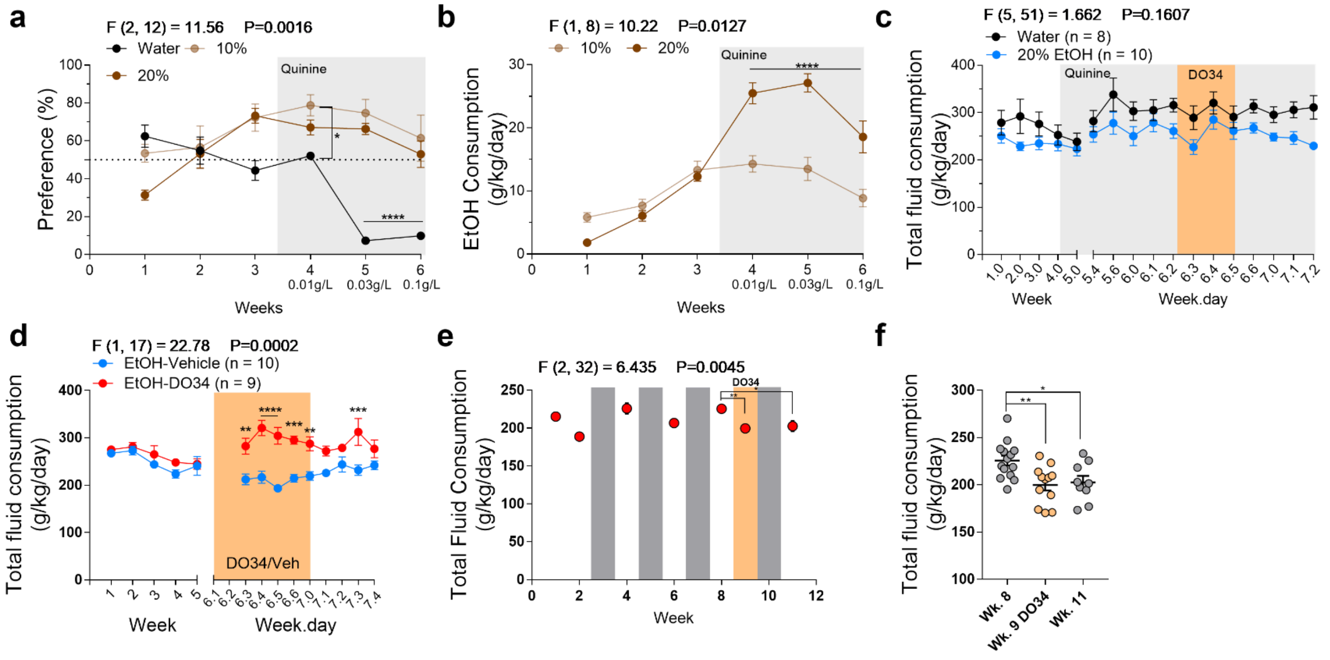


**Figure S7: Mice reliably drink quinine-adulterated EtOH and DO34 has variable but minimal effects on total fluid consumption across EtOH drinking models. (a)** Mice showed significantly higher preference for EtOH + quinine compared to water + quinine at 0.03 and 0.1 g/L quinine. **(b)** Mice drank significantly higher levels of 20% EtOH + quinine compared to 10% EtOH + quinine. **(c)** DO34 treatment had no effect on total fluid consumption in the aversion-resistant drinking model. **(d)** DO34 increased total fluid consumption in the relapse drinking model. **(e)** DO34 treatment reduced total fluid intake in the chronic intermittent ethanol model compared to the baseline week and fluid consumption remained decreased during the recovery week. **(f)** Graph depicting individual mouse total fluid consumption during baseline, treatment, and recovery weeks. Fluid consumption is lower in treatment and recovery weeks compared to baseline, but treatment and recovery consumption levels are not significantly different. **(a-d)** Data were analyzed by repeated measures two-way ANOVA followed by a Holm-Sidak test for multiple comparisons. **(e-f)** Data were analyzed by one-way ANOVA followed by a Holm-Sidak test for multiple comparisons. **(a-b)** Time points during quinine exposure were analyzed. **(c)** Time points 6.2 – 7.0 were analyzed (to include baseline, drug treatment, and two recovery points). **(d)** All time points were analyzed. **(e-f)** Time points 8, 9, and all were analyzed. Multiple comparisons in **e** were referenced to baseline whereas all means were compared in **f.** *P* and *F* values for main effects of drug or quinine treatment reported on graphs. $n$ = 5 mice per group in (**a-b**). $n$ = 9-14 mice in (**e-f**). Sample size $n$ reported on graphs in **(c-d)**. Significance for post-hoc multiple comparisons reported on graphs (*P<.05, **P<0.01, ***P<0.001, ****P<0.0001). Data are mean ± SEM.

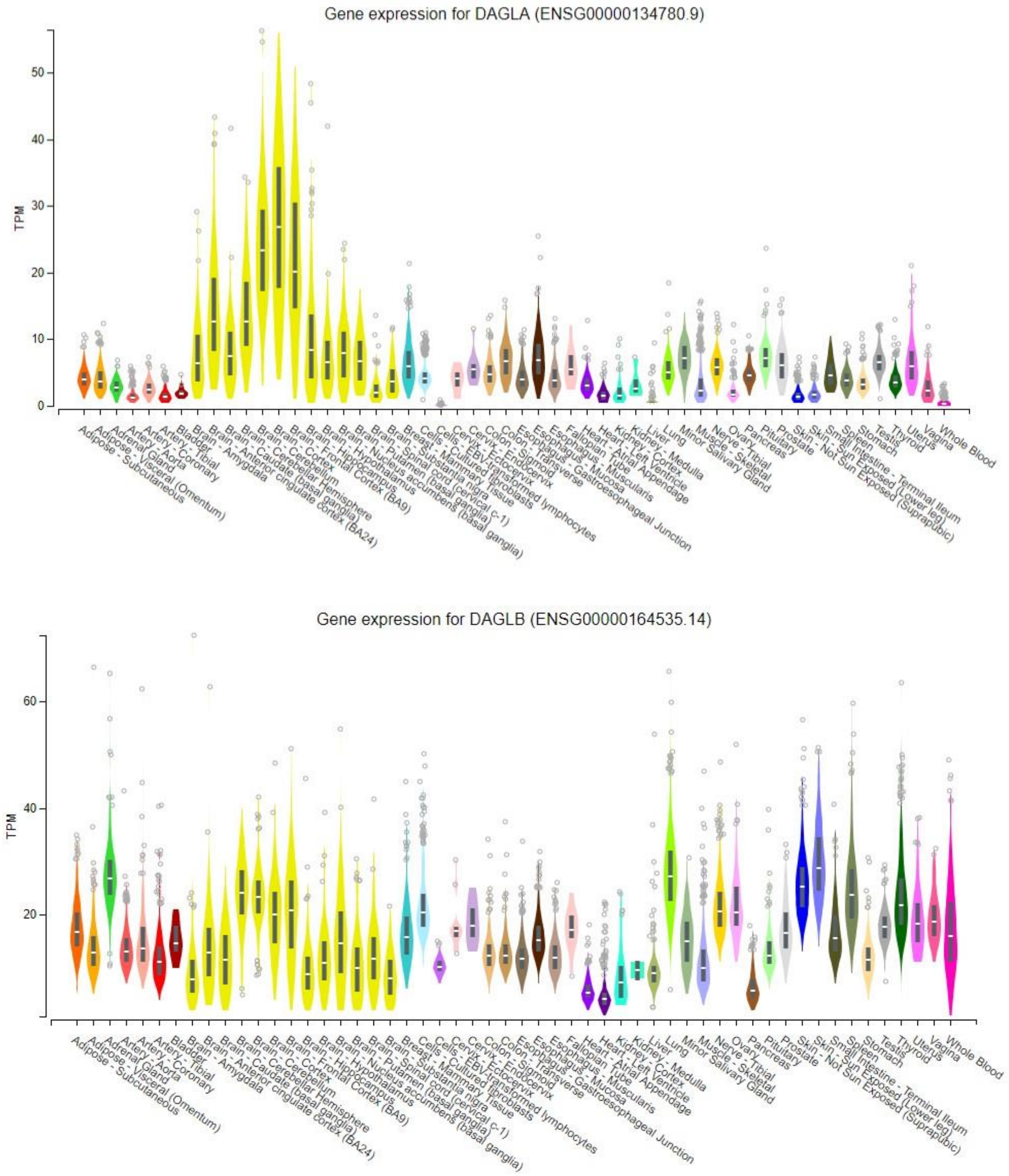


**Figure S8: GTEx data on tissue-specific expression of *DAGLA* and *DAGLB* expression: (top)** Expression profile for *DAGLA*, demonstrating highest expression in brain tissues (yellow). **(bottom)** Expression profile for *DAGLB*, demonstrating ubiquitous tissue distribution. Data are expressed as transcripts per million reads (TPM), and obtained from the GTEx Portal (dbGaP Accession phs000424.v8.p2; see https://gtexportal.org/home)

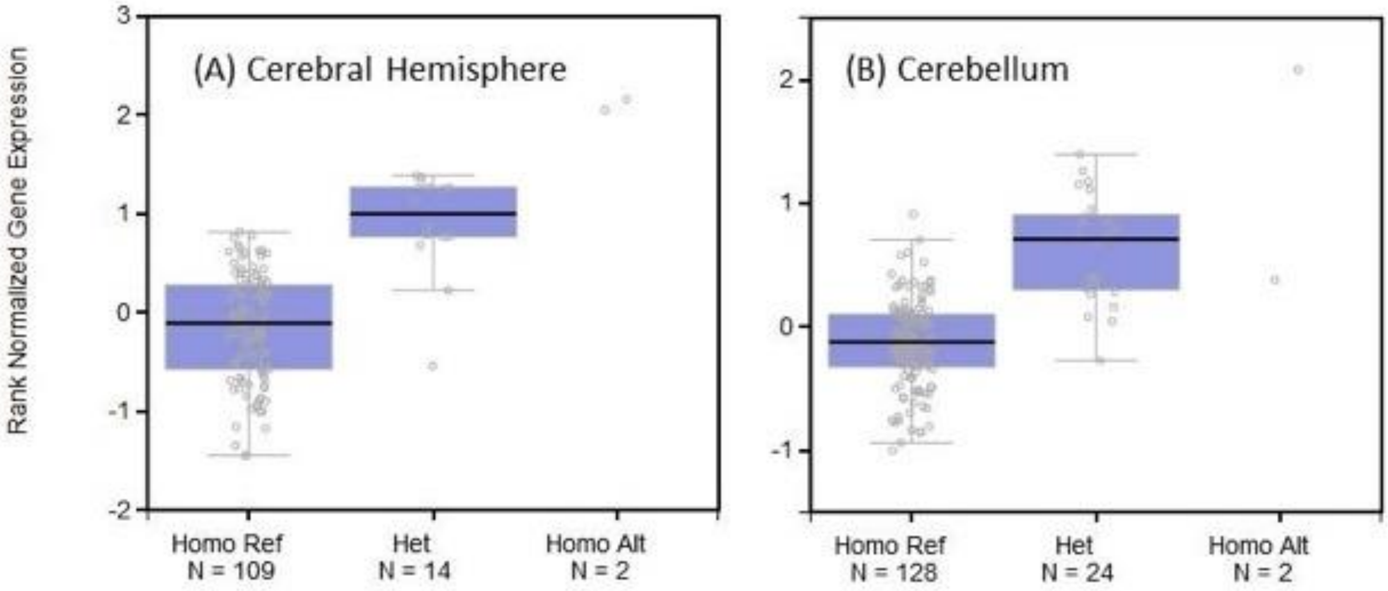


**Figure S9: GTEx data on the effect of *DAGLA* genetic variant rs11604261 on relative gene expression:** The effect of the common variant rs11604261 on DAGLα expression in two brain regions. Normalized expression values are plotted for the three genotypes: homozygous reference, heterozygous, and homozygous alternative allele. Data obtained from the GTEx Portal (dbGaP Accession phs000424.v8.p2; see https://gtexportal.org/home).
